# CycleFlow quantifies cell-cycle heterogeneity *in vivo*

**DOI:** 10.1101/2020.09.10.291088

**Authors:** Adrien Jolly, Ann-Kathrin Fanti, Ines Gräßer, Nils B. Becker, Thomas Höfer

**Affiliations:** Division or Theoretical Systems Biology, German Cancer Research Center (DKFZ), Im Neuenheimer Feld 280, 69120 Heidelberg, Germany; Division of Cellular Immunology, German Cancer Research Center (DKFZ), Im Neuenheimer Feld 280, 69120 Heidelberg, Germany

## Abstract

While the average cell-cycle length in a cell population can be derived from pulse-chase experiments, proliferative heterogeneity has been difficult to quantify. Here we describe CycleFlow, a broadly applicable method that applies Bayesian inference to combined measurements of EdU incorporation and DNA content. CycleFlow accurately quantifies the fraction of proliferating versus quiescent cells and the durations of cell-cycle phases of the proliferating cells *in vitro* and *in vivo*.

## Main

Owing to advances in live-cell microscopy, we are gaining deep understanding of how eukaryotic cells transition between quiescence and proliferation and control the length of their division cycle^1–10^. These studies rely on direct observation of cell cycles *in vitro*. Similar quantification of cell cycles and quiescence during development, tissue renewal and tumor growth in the intact organism has not been possible by imaging, and existing proliferation assays *in vivo* only partially address the key questions. In particular, the rule of thumb that high S and G_2_/M fractions indicate rapid cell proliferation (as measured by DNA staining or single-cell RNA-sequencing (scRNA-Seq) data) can mislead, as we will see below. To quantify cell proliferation, pulse-chase experiments with thymidine analogs or other labels have been widely employed^11^ and, recently, improved by using sophisticated double-labeling techniques^12,13^. However, the quiescent fraction is not resolved in these experiments, which also confounds the estimation of cell-cycle duration. Quiescent cells are often distinguished from cycling cells by (negative) staining for Ki67, but the protein continues to be expressed after cells have stopped cycling, complicating the interpretation of the results^14^. An alternative assay for quiescence, long-term retention of a diluting label, also has limitations, as initially quiescent cells will not be labeled when using thymidine analogs, and dilution of fluorescent histone variants is difficult to distinguish from degradation^15^. Also, there is no generic gene-expression signature for quiescence that could be detected in scRNA-Seq data. Thus, quiescence has largely remained a qualitative rather than quantitative concept despite its key role in stem cell differentiation and therapy resistance.

Here we describe CycleFlow, a time-efficient method that jointly quantifies the quiescent fraction in a cell population and the duration of the cell-cycle phases in the proliferating cells, both *in vitro* and *in vivo*. CycleFlow combines two readily applicable experimental techniques — pulse-chase with a thymidine analog (e.g., 5-ethynyl-2’-deoxyuridine, EdU) and total DNA staining ^16^ — with Bayesian parameter inference. The key idea is to label a cohort of S phase cells with a brief pulse of tymidine analog and then use the DNA content as a coordinate to follow the labeled cells through the cell cycle, eventually observing re-entry of divided cells into the cell cycle or transition into a quiescent state (Fig. 1a). We model label dynamics during cell-cycle progression mathematically and, confronting the model with a time series of flow-cytometry snapshots, infer the fractions of proliferating versus quiescent (G_0_) cells and the time the former spend in the G_1_, S and G_2_/M phases of the cell cycle.

**Fig. 1.**
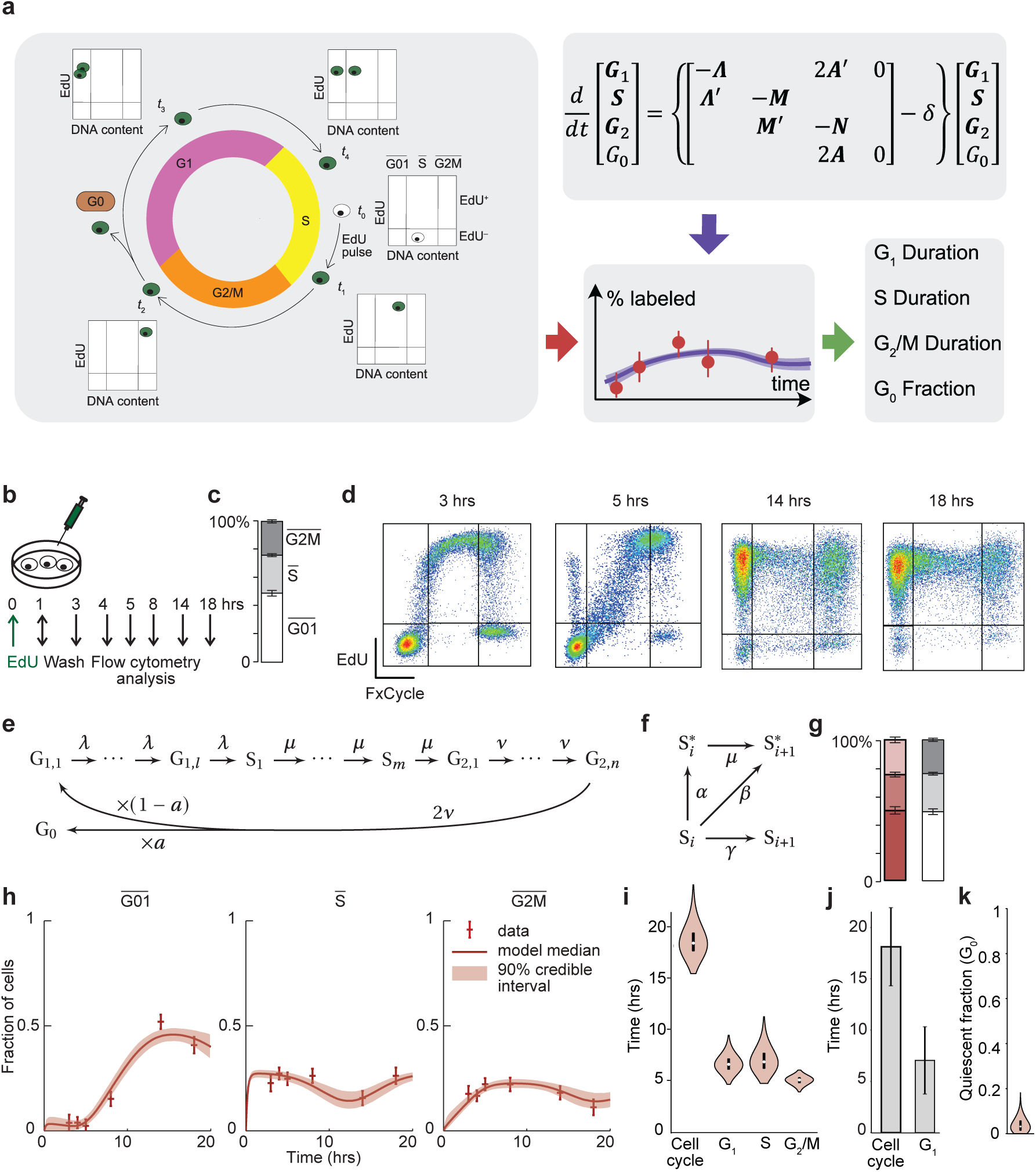
Outline of CycleFlow and application to cancer cell proliferation *in vitro*. **a**, Schematic of CycleFlow: The progression of EdU labeled cells through the cell cycle is tracked over time, the mathematical model is fitted to the data, and cell-cycle parameters are estimated via Bayesian inference. **b**, EdU pulse-chase experiment for cells in culture. **c**, Distribution of cells in the 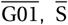, and 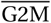 gates as determined by DNA content for TET21N (averaged over all time points). Error bars indicate standard error of the mean (SEM); n = 25. **d**, Progression of EdU labeled TET21N cells through the cell cycle, as defined by DNA content. Four representative flow-cytometry snapshots are shown; mean fractions of cell subpopulations (with SEM) over all experimental repeats at the corresponding times are indicated in **h. e**, Schematic of mathematical model for cell-cycle progression; cell-cycle phases are divided into sub-phases. Parameters: *λ, µ* and *v*, progression rates through subphases of G_1_, S and G_2_/M, respectively, with *l, k* and *n*, denoting subphase numbers; *a*, probability of cell cycle arrest. **f**, Mathematical Model of EdU incorporation during cell progression in S phase. *a* + *β*, total EdU incorporation rate; *β* +*γ* = *µ*, progression rate. **g**, Model fit (left diagram) versus experimental data (right diagram, same as **c**) for the distribution of cells in the 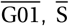, and 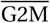 gates. Error bars indicate 90% credible intervals (left diagram). **h**, Time courses of EdU-labeled TET21N cells in 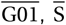, and 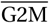 gates compared between experimental data (error bars, pooled SEM; n = 3 to 6 per time point) and model fit. Population sizes are given as fractions of total cells. **i**, Duration of total cell cycle, G_1_, S and G_2_/M phases of TET21N cells inferred by CycleFlow. White dots, median values; black bars, interquartile ranges. **j**, Mean duration of cell-cycle and G_1_ phase of TET21N, expressing the Cdt1 FUCCI degron, measured with time lapse microscopy (data taken from Ref. ^10^). **k**, Quiescent fraction of TET21N cells inferred by CycleFlow.

We first applied CycleFlow to the MYC-driven proliferation of a cancer cell line, TET21N, in culture. This system provides a ground truth for evaluating the performance of CycleFlow, as cell-cycle duration and absence of quiescence have been established by time-lapse imaging^10^. We grew TET21N cells in the exponential phase, added EdU to the culture medium and washed after one hour; cells were collected at six time points over the course of 18 hours, and analyzed by flow cytometry for DNA and EdU content (Fig. 1b). As expected for an exponentially growing population, the proportions of the cell-cycle phases (Fig. 1c) stayed approximately constant; for brevity, we denote the corresponding gates by 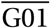 (2n DNA content), 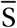 (DNA content intermediate between 2n and 4n) and 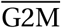 (4n DNA content). Three hours after EdU application, all cells in 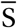 were EdU-positive (Fig. 1d). At 5 hours, cells in the initial stages of 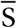 were not EdU-labeled, implying that EdU was no longer available for *de novo* labeling. This allowed us to follow the progression of the cohort of initially EdU-labeled cells around the cell cycle; at 5 hours, these cells were predominantly in late 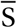 and 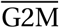. The labeled cohort returned to 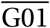 (14 hours), remaining clearly distinguishable from EdU-negative cells. From 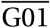, labeled cells re-entered into the cell cycle and progressively appeared again in 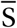 (14 and 18 hours). Thus, EdU pulse labeling allows following a cohort of cells over one cycle.

However, there are several difficulties in interpreting these data in a straightforward manner. First, the cells within the EdU-labeled cohort do not transit synchronously to G_2_, which could be due to variable starting points in S phase at the time of labeling or cell-to-cell variations in the speed of progress. Second, cells within 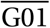 may be either in G_0_ or G_1_. Third, the precise time course of EdU availability is not measured. Finally, the assignment of cell-cycle phases based on DNA content has limited resolution: Note that, at 3 hours, some cells at the right-hand edge of 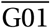 were EdU-positive, implying that these had already entered S phase but were not identified as such by DNA staining (Fig. 1d). Likewise, it is possible that some labeled cells in 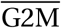 were still in late S phase. To address these difficulties, CycleFlow bases the interpretation of the experimental data on a mathematical model of cell-cycle progression and EdU labeling that takes into account all relevant features (Fig. 1e,f; Supplementary Table 1; Online Methods). Specifically, the model describes the progression of a cell through the cycle as a Markov process, dividing each cycle phase (G_1_, S and G_2_/M) into subphases in order to account for phase length variability^8^ (Fig. 1e). Upon division, cells may continue to cycle or enter quiescence (G_0_). Release from G_0_ may occur but can be neglected on the short timescale of the experiment. Cells may also be lost from any cycle phase or G_0_, e.g., due to cell death or differentiation. As long as EdU is present in the medium (as estimated from the data), cells in S phase can incorporate it and become labeled (Fig. 1f). Finally, the model exploits the joint information on DNA content and EdU labeling available for each cell, rather than DNA-content-based gates only, to assign cells to the cycle phases.

Using Bayesian inference, we estimated the model parameters from the data by Monte-Carlo sampling from the posterior distribution. The model accurately fitted the experimental data within measurement error (Fig. 1g,h), inferring a cycle length of 18.4 (16.7, 21.1) hours (posterior median and (5%, 95%) quantiles) and phase lengths for G_1_, S and G_2_/M of 6.5 (5.5, 8) hours, 6.8 (5.5, 9) hours and 5 (4.3, 5.6) hours, respectively (Fig. 1i). Using direct observation by time-lapse microscopy of TET21N cells expressing the FUCCI Cdt1 degron^10^, reliable values were obtained for total cycle and G_1_ phase lengths, and these matched the results of CycleFlow (Fig. 1j). Moreover, CycleFlow inferred the fraction of G_0_ cells to be negligible (Fig. 1k), as observed^10^. Taken together, these findings validate CycleFlow.

Next, we applied CycleFlow to a developmental process *in vivo*, where the extent of quiescence has been controversial—the differentiation of T cells in the thymus. We focused on the CD4^+^ CD8^+^ double-positive (DP) thymocytes, the development of which is accompanied by a transition from proliferation to quiescence^17^. Varying estimates of the quiescent fraction (30-90%) have been put forward but direct quantification has been lacking^18^. To address this question, we injected a single intra-peritoneal dose of EdU into young adult mice and performed EdU and DNA staining at consecutive time points (Fig. 2a). Gating by DNA content showed >90% of DP thymocyte in 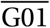 (Fig. 2b). We observed rapid termination of EdU labeling (Fig. 2c, 3 hours) and, as before, return of the labeled cohort to 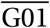 (Fig. 2c, 3 and 5 hours) followed by re-entry into 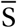 and 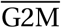 (Fig. 2c, 14 hours). Of note, a large fraction of cells remained unlabeled. All these features were captured by the mathematical model, which reproduced both the steady-state cell-cycle distribution (Fig. 2d) and the dynamics of labeled cells (Fig. 2e).

**Fig. 2.**
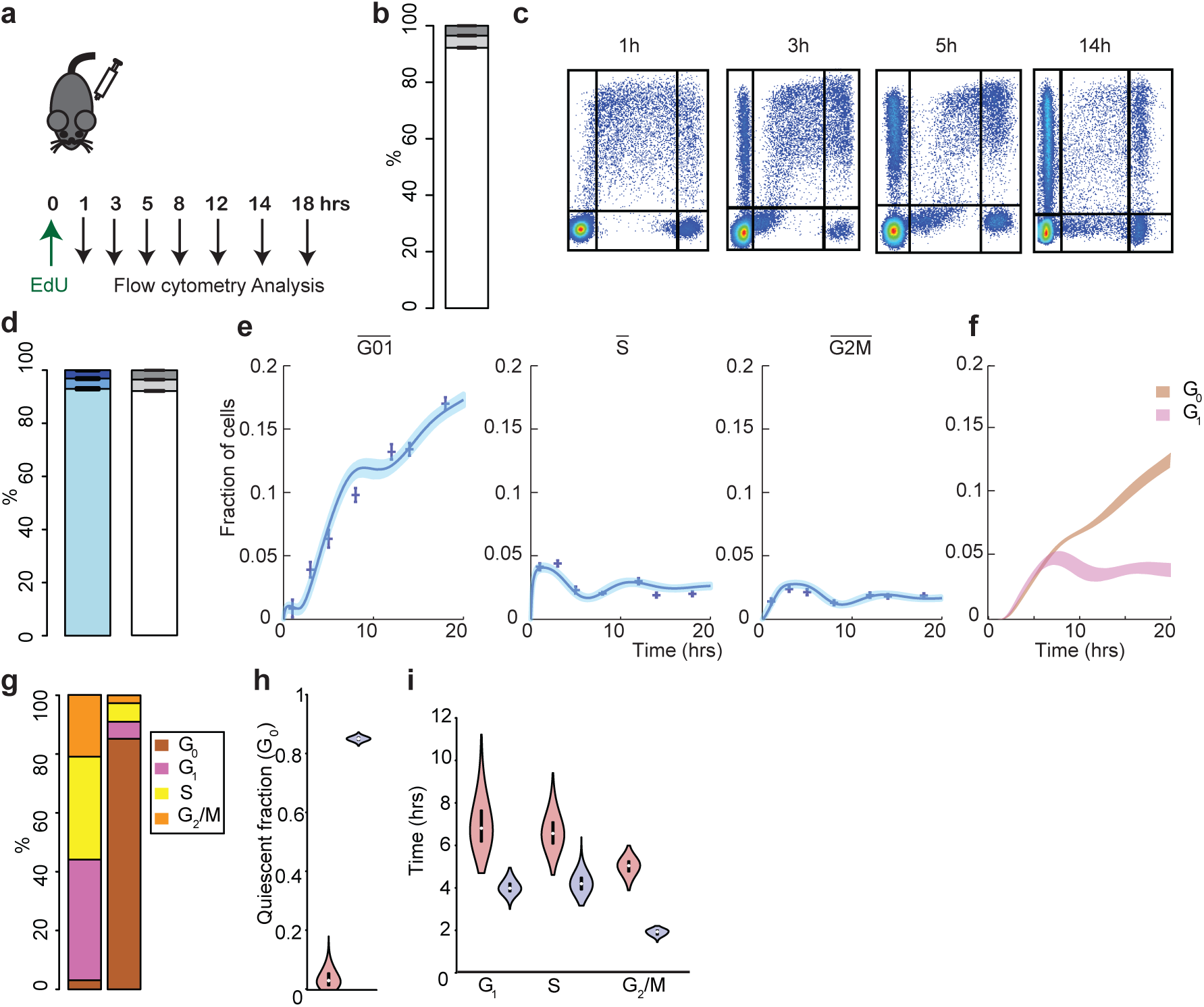
Application of CycleFlow to determine quiescence and proliferation rate of DP thymocytes *in vivo*. **a**, Schematic of the EdU pulse-chase experiment. **b**, Distribution of cells in the 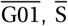, and 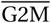 gates as determined by DNA content for DP (averaged over all time points). Errors bars indicate standard error of the mean (SEM); n = 31. **c**, Progression of EdU-labeled DP thymocytes through the cell cycle, as defined by DNA content. Four representative flow-cytometry snapshots are shown; mean fractions of cell subpopulations (with SEM) over all experimental repeats at the corresponding times are indicated in **e. d**, Model fit of the distribution of cells in the 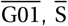, and 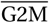 gates, error bars indicate 90% credible interval (left bar) and corresponding data (right bar; same as **b**). **e**, Time courses of EdU-labeled DP cells in 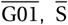, and 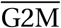 gates compared between experimental data (error bars, pooled SEM; n = 3 to 6 per time point) and model fit. Population sizes are given as fractions of total cells. **f**, Inferred fractions of EdU-labeled G_0_ and G_1_ cells in the 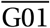 gate. **g**, Inferred distribution of cell-cycle phases including G_0_ (median values). Left bar, TET21N cells; right bar, DP thymocytes. **h**, Inferred fraction of quiescent cells in TET21N cells (left) and DP thymocytes (right). White dots, median values; black bars, interquartile ranges. **i**, Duration of G_1_, S and G_2_/M phases of DP thymocytes (blue) compared to TET21N cells (red) inferred by CycleFlow. White dots, median values; black bars, interquartile ranges.

Interestingly, although the fraction of labeled cells in 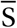 and 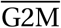 plateaued at low values already after a few hours, the labeled fraction in 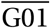 kept increasing for the duration of the experiment (18 hrs), raising the question of which process fuels the continuing label accumulation. 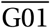 contains cells in G_1_ and G_0_; the model separates these contributions and shows that, upon exit from mitosis, nearly equal fractions of cells enter quiescence and continue cycling (Fig. 2f). While the fraction of labeled cycling cells quickly plateaus (similar to the TET21N cells, Supplementary Fig. 1), the labeled quiescent cells continue to accumulate. Thus, in DP thymocytes proliferation precedes quiescence, which is consistent with the known developmental progression^18^.

Comparing the experimental data and the inferred cell-cycle parameters for TET21N cells and DP thymocytes illustrates how the simple rule of thumb of judging proliferation rate by S and G_2_/M fractions can fail. The cell-cycle phase distributions show much higher 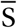 and 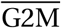 fractions for TET21N cells (compare Figs 1c and 2b), which would indicate a higher proliferation rate in this population compared to the DP thymocytes. However, the proliferating DP thymocytes actually cycle faster than TET21N cells, but this is masked by the high fraction of quiescent thymocytes. CycleFlow allows us to see beyond population averages and disentangle cell-cycle heterogeneity by inferring the underlying cell-cycle phase distribution (Fig. 2g), rather than observing gated fractions. Specifically, TET21N cells proliferate uniformly (quiescent fraction 3 (0, 9) %), with 18.4 (16.7, 21.1) hours average cycle length (cf. Fig. 1i,k). By contrast, the large 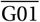 fraction in DP thymocytes consists mainly of quiescent cells, which constitute 85 (84, 86) % of all cells (Fig. 2h), corroborating the existence of a large quiescent fraction^18^. The remaining 15 (14, 16) % of cells cycle extremely rapidly, with a cycle length of 10 (9.4, 10.9) hours (Fig. 2i). We note that, whereas Chao et al. (Ref 8) found that cycle length of different cell types was primarily set by the duration of G_1_, all phases appear to be shortened in DP thymocytes (Fig. 2i), with a particularly swift passage through G_2_/M. This finding indicates that proliferation rate can be regulated at all stages of the cell cycle^19^.

To conclude, the main advance of CycleFlow is that it disentangles quiescent (G_0_) from cycling (G_1_) cells with 2*n* DNA content. In turn, this enables determination of the true cell-cycle phase durations of the proliferating cells. In so doing, our mathematical framework overcomes salient problems of previous approaches, including the imperfect overlap between cell-cycle phases and DNA content gates and the gradual removal of EdU. CycleFlow is equally applicable for steady-state (e.g., thymocytes) and expanding (e.g., TET21N cells) cells, both *in vivo* and *in vitro*. Application to other cell types is straightforward: the measurements should extend over roughly one cell cycle with several (typically 4-8) loosely distributed time points.

## Methods

### Experimental

#### Labeling and analysis of TET21N cells *in vitro*

SH-EP TET21/N (TET21/N) cells were grown in RPMI 1640 medium supplemented with 10% Fetal Calf Serum (FCS) at 37°C, 5% CO_2_ and 88% humidity. For each sample, 1.5 × 10^6^ cells were seeded on 15 cm Petri dishes one day before EdU treatment and then treated with EdU (Invitrogen) at a final concentration of 10 µM in the culture medium. Cells were then harvested and fixed in 4% paraformaldehyde solution in PBS and kept in 90% methanol 10% PBS solution at −20°C. For flow cytometry analysis, cells were washed in washing buffer (1% Bovine Serum Albumin(BSA), 0.1% TritonX in PBS) and resuspended in PBS supplemented with 1% BSA. Cells were then permeabilized and the Click-it reaction was performed using the Click-iT Plus EdU Alexa Fluor 488 flow cytometry Kit (Invitrogen) according to the manufacturer’s protocol. For total DNA staining, cells were resuspended in FxCycle™ Violet (Thermofisher Scientific) solution (1:1000 in Click-it Permeabilization buffer) prior to flow cytometry measurement. Data were acquired on a MACSQuant VYB (Miltenyi Biotec) and cell populations were analyzed with FlowJo 10.

#### Mice

C57BL/6J mice were used, both female and male. Mice were kept in individually ventilated cages under specific pathogen-free conditions in the animal facility at the German Cancer Research Center (DKFZ, Heidelberg). All animal experiments were performed in accordance with institutional and governmental regulations, and were approved by the Regierungspräsidium (Karlsruhe, Germany).

#### Labeling of thymocytes *in vivo* and analysis

Mice of ages between eleven and sixteen weeks were injected intraperitoneally with 1 mg EdU (Invitrogen) diluted in sterile PBS. At several time points after injection mice were sacrificed and thymi harvested. Thymi were mashed in a 40 µm filter with the plunger of a syringe. To identify dead cells, cells were incubated in Zombie Red™ Fixable Viability dye (Biolegend) solution (1:1000 in PBS). Fc receptors were blocked by incubating cells in PBS supplemented with 5% FCS with 250 µ g.ml^−1^ purified mouse IgG (Jackson ImmunoResearch Laboratories). Antibody stainings were performed in PBS/5% FCS on ice for 30 minutes with optimal dilutions of commercially-prepared antibodies. The following antibodies were used: CD8a allophycocyanin (APC) (53-6.7), TCR gd BV421 (GL3) from eBioscience; CD4 phycoerythrin (PE) (H129.19), Ter119 BV421 (Ter119) from BD Pharmingen; CD11b BV421 (M1/70), CD19 BV421 (6D5), NK1.1 BV421 (PK136), Gr-1 BV421 (RB6-8C5) from Biolegend. The lineage cocktail was composed of CD11b, CD19, Ter119, NK1.1, Gr-1 and TCR gd. After antibody staining, cells were fixed, permeabilized and the Click-it reaction was performed using the Click-iT Plus EdU Alexa Fluor 488 flow cytometry kit (Invitrogen). For total DNA staining, cells were resuspended in FxCycle™ Violet solution (1:1000 in Click-it Permeabilization buffer) prior to flow cytometry measurement. Data was acquired on a BD LSRFortessa™ cell analyzer (Becton Dickinson) and cell populations were analyzed with FlowJo 10. Double-positive thymocytes were defined as lineage^−^ CD4^+^CD8^+^.

### Mathematical inference

#### Model for cell-cycle progression

Within a growing cell population, each cell progresses through the cycle phases G_1_, S and G_2_ (Fig. 1e). Each phase requires a stochastic time to complete. We capture this variability by dividing the cycle phases into subphases: G_1,1_ through G_1,*/*_, S_1_ through S_m_ and G_2,1_ through G_2,*n*_, respectively. Each subphase corresponds to an approximate progress within the respective cycle phase but has no further biological meaning. We then model the transitions from one subphase to the next (and from the last subphase of a cycle phase to the first subphase of the next) as rate processes with rates *λ, µ* or *v*, for G_1_, S and G_2_, respectively. In this model, the times to complete each of the cycle phases are Erlang-distributed random variables, with means 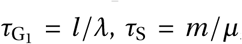, and 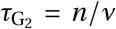, and coefficients of variation (CVs), 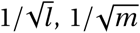 and 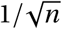, respectively. At the end of the cycle, a cell divides to produce two daughters. A daughter cell may then, with probability *a*, exit the cell cycle into the arrested state G_0_, or, with probability 1 − *a*, restart the cycle in phase G_1,._ We consider entry into G_0_ to be irreversible on the time scale of the experiment, and therefore exclude any transition from G_0_ back into the cycle. Finally, cells may be lost from any cycle phase or G_0_ with rate *δ*, due to cell death or differentiation.

The dynamics of subpopulations is then described by a linear system of ordinary differential equations. Written in matrix form,

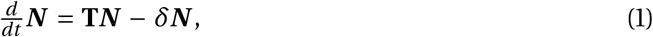

where the vector of cell numbers in the various cycle subphases is defined as

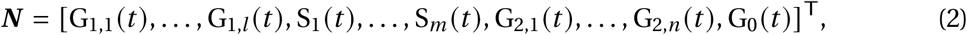

and the rate matrix

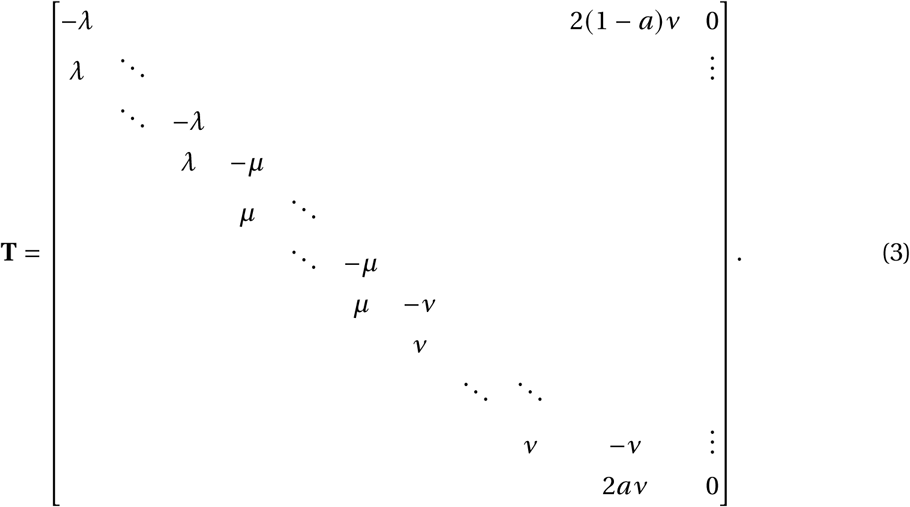

In a cell population deep in the exponential growth phase, the total cell number *N* (*t*) increases exponentially, while the proportions of cells in the various subphases are constant; we call this regime *steady growth*. At steady growth,

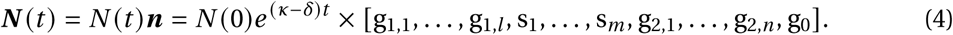

Here, we have denoted the population growth rate by (*κ* − *δ*). The proportions of cells in the subphases are given by the steady-growth distribution ***n***; denoting the constant vector of ones by **1** = [1, *…*, 1]^T^, the normalization of ***n*** reads **1**^T^***n*** = 1. Eq. 1 then yields

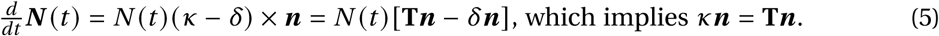

Thus *κ* is the dominant eigenvalue of the matrix **T**, and ***n*** is the associated normalized eigen-vector. The eigenvalue *κ* can be interpreted as the population growth rate in the absence of loss.

#### Steady-growth distribution

To find ***n***, we start by normalizing Eq. 5:

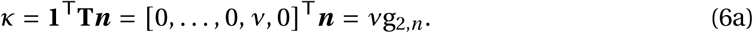

Eq. 6a implies that the total rate of production of new cells *κN* equals the population in the final subphase G_2,*n*_ times its rate of completion. Now consider steady growth with production of new cells, where G_2,*n*_, *κ* > 0. Starting from Eq. 6a and backsubstituting using Eq. 5, we compute the remainder of the cycle subphase distribution in steady growth as a function of *κ*:

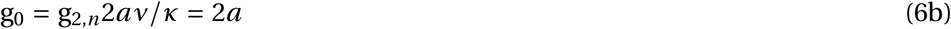

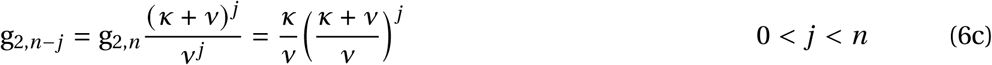

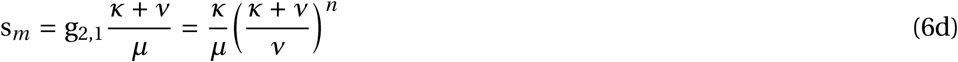

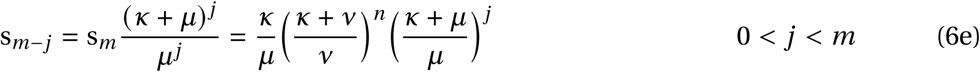

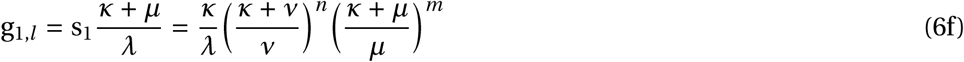

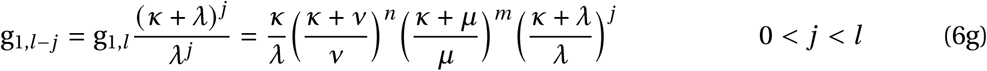

To complete the calculation, the dominant eigenvalue *κ* can be found numerically by solving Eq. 5 for given transition rates and subphase numbers. For steady growth to be possible, *a* ≤ 1/2 is required, since otherwise each cell has less than one proliferating daughter cell on average.

#### Steady-state distribution

To capture cell populations that are maintained at a fixed size within the model Eq. 1, we consider the loss rate *δ* to be subject to an implicit homeostatic regulation. This regulation maintains *δ* ≡ *κ*, so that proliferation and loss balance, and no net growth occurs. In this way, homeostasis is treated as a marginal case of steady growth, Eq. 4 with vanishing effective growth rate, so that the subphase populations ***N*** = *N****n*** are constant in time. In particular, we can compute the steady-state distribution ***n*** by Eqs. 4 with the replacement *κ* → *δ* everywhere. If desired, one also obtains the homeostatic set point of *δ*, by solving the eigenvalue problem Eq. 5, with *κ* → *δ*.

#### Kinetics of labeling

In order to relate the cycling model Eq. 1 to experimental data, we include the dynamics of label incorporation and inheritance. As long as EdU label is available, unlabeled cells in S phase incorporate it, thereby transitioning into a labeled state. The labeled states (denoted S^*^, 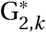 etc.) are defined operationally: Those cells that are gated above the background fluorescence level in the EdU fluorescence channel are considered labeled. In practice we then find that transition to the labeled state requires only a small part of S phase, in other words, the rate of label acquisition *ϵ* > *µ*/*m*. However it’s unclear if label acquisition is also faster than progress from one S subphases to the next, *ϵ* ≷ *µ*. Therefore, we consider S phase progress and labeling to be parallel processes, see Fig. 1f. In this scheme, the three new rates *α, β* and *γ* are constrained by

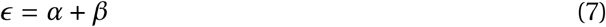

setting the total rate of labeling to be *ϵ*, and

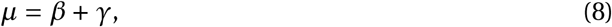

which states that the total progress rate of unlabeled cells through S phase is undisturbed by label incorporation. We impose a third constraint for convenience,

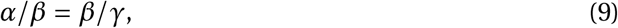

which then uniquely determines the three rates for given *E*:

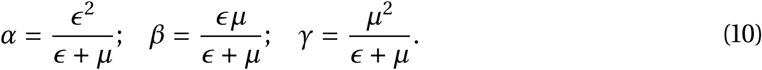

Since the chase phase of the experiment remains shorter than two cell cycles, we do not observe nor include in the model any delabeling by dilution of EdU.

#### Labeled population dynamics

Finally, we collect the previous expressions into a system of linear ODEs for the dynamics of all, labeled and unlabeled subpopulations. The vector of average labeled cell numbers is

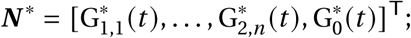

unlabeled cell numbers are denoted by ***N***°. They add up to the total cell numbers ***N*** = ***N***°+***N***^*^.

We require that steady growth or steady state has been attained before labeling begins. Thus up to the label application at *t* = 0, ***N***° = ***N*** follows Eq. 4, with steady-growth distribution ***n*** satisfying Eq. 5. Labeling transfers cells into the labeled populations, so that at times *t* > 0, Eq. 1 holds for ***N*** but not for ***N***° nor ***N***^*^.

To describe the dynamics of the labeled population ***N***^*^, we encode the labeling kinetics in matrix form:

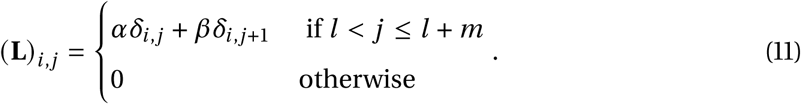

The labeling matrix **L** has the same dimensions as **T**; its nonzero entries are *α* on the diagonal for all subphases of S, and *β* below any *α*. Here, *α* and *β* are given by Eqs. 10. It can be verified straightforwardly that **L*N***° is the flux from unlabeled to labeled compartments as shown in Fig. 1f. Using this notation,

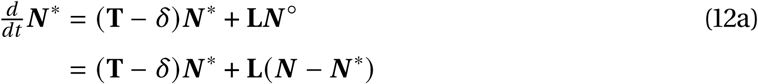

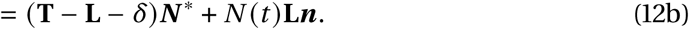

In Eq. 12a, we have used the fact that already-labeled cells follow the undisturbed cycle progression (**T** − *δ*); in Eq. 12b, we have inserted the undisturbed steady-growth expansion of the total population. Finally, we rewrite this system in terms of the dynamics of the (time-dependent, non-normalized) labeled fractions, 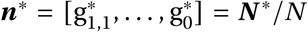. From Eq. 12b and using exponential growth of *N* (*t*), we obtain

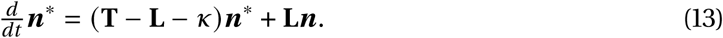

Importantly, in experiments EdU supply is stopped at the end of an initial labeling pulse, and thereafter, the availability of free EdU decreases gradually as it is consumed by cells or otherwise degraded. We model this decrease by an exponential decrease of the total labeling rate

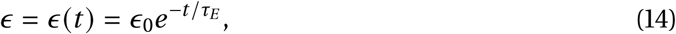

where *τ*_*E*_ is an adjustable parameter. In this way *α, β* (Eq. 10) and therefore **L** (Eq. 11) become time-dependent.

#### Parameter estimation

Experimentally, cycle phase is assigned by DNA content. Thus, cells in G_0_ or G_1_ fall into a single experimental gate, 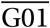. Although most DNA-replicating cells are correctly counted in the corresponding 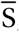 gate, the gating procedure inevitably assigns some cells that have started DNA replication to the 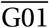 gate, and some cells that have completed replication to the 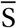 gate rather than the 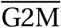 gate. When deriving model predictions, we account for this crosstalk by assigning the first 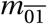 subphases of S phase to 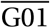, and the last 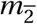 subphases to 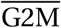, so that only the innermost 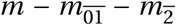 subphases S_*i*_ are assigned to the gate 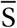, as shown in Eq. 15.

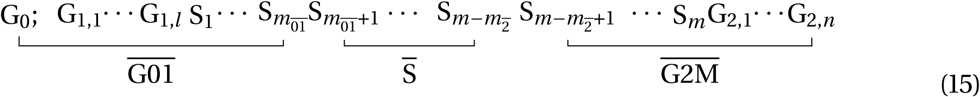

In order to compute model predictions, we first solve Eq. 5 for the steady-growth distribution ***n***. From ***n*** we obtain the time-independent total cell fractions in each experimental gate,

*e*.*g*.

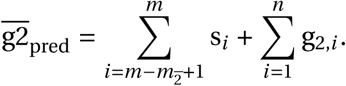

We then solve system Eq. 13 numerically, for the initial condition ***n***^*^(0) = **0**. From ***n***^*^(*t*), we obtain predictions for the time-dependent labeled fractions, *e*.*g*.

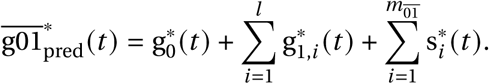

In order to then compare model predictions with experiments, we evaluate the negative log-likelihood function in a standard way, as

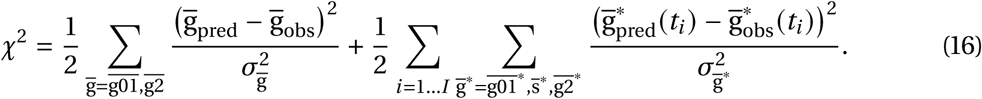

Here, the first sum collects time independent terms; the experimental observations 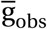 are averages over all time-points and experimental repeats, and the associated uncertainties 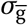 are standard errors of these averages. Only two gates are included, since normalization fixes the third. The second sum runs over the *I* experimental time points and is not constrained by normalization. The corresponding uncertainties are estimated as standard errors over experimental repeats of the mean labeled gate fractions; for robustness, we pool these errors and assign the same 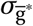 for all time points.

Based on the log-likelihood Eq. 16, we then perform Markov Chain Monte Carlo sampling to evaluate the posterior distribution over model parameters. The full set of model parameters and the allowed ranges of their uniform prior distributions are given in Supplementary Table 2. The credible intervals resulting from sampled posterior distributions for DP and TET21N are shown in Supplementary Table 3.

### Implementation and Software

The model was simulated using the Python programming language with the package Scipy v.0.13.0 and Monte Carlo Markov Chain (MCMC) sampling was performed with the package Emcee v.3.02 with default settings. In accordance with the program’s documentation, a chain was deemed to have converged when the autocorrelation times for every parameters exceeded 50 times the length of the chain. Results of the MCMC sampling were then processed with Matlab R2013B (MathWorks) and R to produce graphical representations.

## Acknowledgments

Funding through the Computational Life Sciences program of the Federal Ministry for Education and Research (FKZ 031L0170A, to TH) and DKFZ core funding are gratefully acknowledged. We thank Katrin Busch and all members of the Höfer group for discussions.

## Contributions

T.H., A.J. and N.B. conceived the Study; N.B. and T.H. supervised the project; A.J and A-K.S. conceived and conducted the experiments on mice; A.J and I.G. conceived and conducted the experiments on TET21N; A.J. and N.B. developed the mathematical model; A.J. implemented the model and A.J. and N.B. analyzed the data; T.H., N.B. and A.J wrote the manuscript and all the coauthors contributed to the manuscript.

## Data availability

All raw Data and codes are available upon reasonable request to the authors.

