## Supplemental Information for "CycleFlow quantifies cell-cycle heterogeneity *in vivo*"

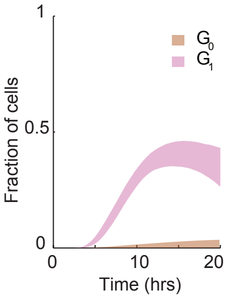


**Supplementary Figure 1 | Inferred time course of EdU-labeled $G_0$ and $G_1$ TET21N cells in the $\overline{\mathrm{G01}}$ gate.** Population sizes are given as fractions of total cells.

| Challenge | Addressed by |
|---|---|
| Cells labeled at any stage of S phase. | Subphases track progress within cycle phases. |
| Variable cycle progression speed. | Distributions for phase durations. |
| Ambiguity between $G_0$ and $G_1$. | Model states are unambiguous cycle phases, not gates. |
| EdU is consumed gradually. | Decrease in labeling efficiency is fitted from data. |
| DNA content reflects cycle phase with limited accuracy. | Cells at boundaries of DNA staining gates assigned to cycle phases via model fit to data. |

**Supplementary Table 1 | Model features introduced to address the main challenges in the interpretation of thymidine analog labeling.**

| Parameter | Description | Unit | Prior range |
|---|---|---|---|
| $\lambda$ | $G_1$ subphase progress rate | $h^{-1}$ | $[0.01, 10]$ |
| $\mu$ | S subphase progress rate | $h^{-1}$ | $[1, 5]$ |
| $\nu$ | $G_2$ subphase progress rate | $h^{-1}$ | $[1, 15]$ |
| $\delta$ | loss rate | $h^{-1}$ | $[0.001, 0.05]$ |
| $a$ | Cycle arrest probability | | $[0, 1]$ |
| $\tau_E$ | EdU degradation time | h | $[1, 5]$ |
| $\epsilon_0$ | initial labeling rate | $h^{-1}$ | 5, fixed |
| $l$ | $G_1$ subphase number | | $[2, 30]$ |
| $m$ | S subphase number | | 15, fixed |
| $n$ | $G_2$ subphase number | | 15, fixed |
| $m_{\overline{01}}$ | S subphases assigned to $\overline{\text{G01}}$ | | $[0, 5]$ |
| $m_{\overline{2}}$ | S subphases assigned to $\overline{\text{G2M}}$ | | $[0, 5]$ |

**Supplementary Table 2 | Model parameters with allowed ranges.**

| Parameter | Description | Unit | TET21N | DP |
|---|---|---|---|---|
| $\lambda$ | $G_1$ subphase progress rate | $h^{-1}$ | $3.5(1.5, 4.9)$ | $0.47(0.4, 0.56)$ |
| $\mu$ | S subphase progress rate | $h^{-1}$ | $2.2(1.6, 2.7)$ | $3.8(3.3, 4.2)$ |
| $\nu$ | $G_2$ subphase progress rate | $h^{-1}$ | $3(2.7, 3.5)$ | $7.7(7.2, 8.8)$ |
| $\delta$ | loss rate | $h^{-1}$ | n.a | $0.013(0.012, 0.014)$ |
| $a$ | Cycle arrest probability | | $0.015(0.0015, 0.045)$ | $0.425(0.42, 0.43)$ |
| $\tau_E$ | EdU degradation time | h | $2.9(2.3, 3.8)$ | $1.1(1, 1.2)$ |
| $l$ | $G_1$ subphase number | | $23(10, 30)$ | $2(2, 2)$ |
| $m_{\overline{01}}$ | S subphases assigned to $\overline{\text{G01}}$ | | $2(0, 3)$ | $4(3, 4)$ |
| $m_{\overline{2}}$ | S subphases assigned to $\overline{\text{G2M}}$ | | $1(0, 2)$ | $0(0, 1)$ |
| $l/\lambda$ | $G_1$ duration | h | $6.5(5.5, 8)$ | $4.2(3.6, 4.9)$ |
| $m/\mu$ | S duration | h | $6.8(5.5, 9)$ | $4(3.6, 4.6)$ |
| $n/\nu$ | $G_2$ duration | h | $5(4.3, 5.8)$ | $1.9(1.7, 2)$ |
| $a/2$ | $G_0$ fraction | | $0.03(0, 0.09)$ | $0.85(0.84, 0.86)$ |

**Supplementary Table 3 | Model parameters medians and 90% Credible Intervals.**

| Parameter | Description | Unit | TET21N | DP |
|---|---|---|---|---|
| $\lambda$ | $G_1$ subphase progress rate | $h^{-1}$ | $3.5(1.5, 4.9)$ | $0.47(0.4, 0.56)$ |
| $\mu$ | S subphase progress rate | $h^{-1}$ | $2.2(1.6, 2.7)$ | $3.8(3.3, 4.2)$ |
| $\nu$ | $G_2$ subphase progress rate | $h^{-1}$ | $3(2.7, 3.5)$ | $7.7(7.2, 8.8)$ |
| $\delta$ | loss rate | $h^{-1}$ | n.a | $0.013(0.012, 0.014)$ |
| $a$ | Cycle arrest probability | | $0.015(0.0015, 0.045)$ | $0.425(0.42, 0.43)$ |
| $\tau_E$ | EdU degradation time | h | $2.9(2.3, 3.8)$ | $1.1(1, 1.2)$ |
| $l$ | $G_1$ subphase number | | $23(10, 30)$ | $2(2, 2)$ |
| $m_{\overline{01}}$ | S subphases assigned to $\overline{\mathrm{G01}}$ | | $2(0, 3)$ | $4(3, 4)$ |
| $m_{\overline{2}}$ | S subphases assigned to $\overline{\mathrm{G2M}}$ | | $1(0, 2)$ | $0(0, 1)$ |
| $l/\lambda$ | $G_1$ duration | h | $6.5(5.5, 8)$ | $4.2(3.6, 4.9)$ |
| $m/\mu$ | S duration | h | $6.8(5.5, 9)$ | $4(3.6, 4.6)$ |
| $n/\nu$ | $G_2$ duration | h | $5(4.3, 5.8)$ | $1.9(1.7, 2)$ |
| $a/2$ | $G_0$ fraction | | $0.03(0, 0.09)$ | $0.85(0.84, 0.86)$ |

**Supplementary Table 4 | Model parameters medians and 90% Credible Intervals.**

| Parameter | Description | Unit | TET21N | DP |
|---|---|---|---|---|
| $\lambda$ | $G_1$ subphase progress rate | $h^{-1}$ | $3.5(1.5, 4.9)$ | $0.47(0.4, 0.56)$ |
| $\mu$ | S subphase progress rate | $h^{-1}$ | $2.2(1.6, 2.7)$ | $3.8(3.3, 4.2)$ |
| $\nu$ | $G_2$ subphase progress rate | $h^{-1}$ | $3(2.7, 3.5)$ | $7.7(7.2, 8.8)$ |
| $\delta$ | loss rate | $h^{-1}$ | n.a | $0.013(0.012, 0.014)$ |
| $a$ | Cycle arrest probability | | $0.015(0.0015, 0.045)$ | $0.425(0.42, 0.43)$ |
| $\tau_E$ | EdU degradation time | h | $2.9(2.3, 3.8)$ | $1.1(1, 1.2)$ |
| $l$ | $G_1$ subphase number | | $23(10, 30)$ | $2(2, 2)$ |
| $m_{\overline{01}}$ | S subphases assigned to $\overline{\mathrm{G01}}$ | | $2(0, 3)$ | $4(3, 4)$ |
| $m_{\overline{2}}$ | S subphases assigned to $\overline{\mathrm{G2M}}$ | | $1(0, 2)$ | $0(0, 1)$ |
| $l/\lambda$ | $G_1$ duration | h | $6.5(5.5, 8)$ | $4.2(3.6, 4.9)$ |
| $m/\mu$ | S duration | h | $6.8(5.5, 9)$ | $4(3.6, 4.6)$ |
| $n/\nu$ | $G_2$ duration | h | $5(4.3, 5.8)$ | $1.9(1.7, 2)$ |
| $a/2$ | $G_0$ fraction | | $0.03(0, 0.09)$ | $0.85(0.84, 0.86)$ |

**Supplementary Table 5 | Model parameters medians and 90% Credible Intervals.**
